# Integration of absolute multi-omics reveals translational and metabolic interplay in mixed-kingdom microbiomes

**DOI:** 10.1101/857599

**Authors:** F. Delogu, B.J. Kunath, M.Ø. Arntzen, T.R. Hvidsten, P.B. Pope

**Author notes:** Corresponding authors: Phillip B. Pope, Francesco Delogu.

## Abstract

Microbiology is founded on well-known model organisms. For example, the majority of our fundamental knowledge regarding the quantitative levels of DNA, RNA, and protein backdates to keystone pure culture-based studies. Nowadays, meta-omic approaches allow us to directly access the molecules that constitute microbes and microbial communities, however due to a lack of absolute measurements, many original culture-derived “microbiology statutes” have not been updated or adapted to more complex microbiome settings. Within a cellulose-degrading and methanogenic consortium, we temporally measured genome-centric absolute RNA and protein levels per gene, and obtained a protein-to-RNA ratio of 10^2^-10^4^ for bacterial populations, whereas Archaeal RNA/protein dynamics (10^3^-10^5^: *Methanothermobacter thermoautotrophicus*) were more comparable to Eukaryotic representatives humans and yeast. The linearity between transcriptome and proteome had a population-specific change over time, highlighting a minimal subset of four functional carriers (cellulose degrader, fermenter, syntrophic acetate-oxidizer and methanogen) that coordinated their respective metabolisms, cumulating in the overarching community phenotype of converting polysaccharides to methane. Our findings show that upgrading multi-omic toolkits with traditional absolute measurements unlocks the scaling of core biological questions to dynamic and complex microbiomes, creating a deeper insight into inter-organismal relationships that drive the greater community function.

## Introduction

The foundations of microbiology have been built within the constrained framework of pure culture studies of model organisms that are grown under controlled steady state conditions. However, we are constantly told that microorganisms grown in artificial isolate conditions behave in a different manner than what they do in a more natural community setting. For example, when *Escherichia coli* is grown axenically in steady state, we can expect that each RNA molecule results in 10^2^ to 10^4^ of the corresponding protein (protein-to-RNA ratio) and the variation in the level of cellular RNA explains ~29% of the variation in the amount of detectable protein^1^. Yet does this notion hold true when a given bacterial population is part of a larger community and subject to transitions from one state of equilibrium to another due to limiting and/or confronting environmental factors? In this context, the exploration of temporal interplay between populations with different lifestyles (comprising metabolism, motility, sporulation, etc.) becomes of primary importance to interpret the changes in fundamental quantities in a microbial community, such as the protein-to-RNA ratio that ultimately impacts the overarching community phenotype(s). In order to perform studies of such design and test if previously defined quantitative data about the functioning of microbes (i.e. protein-to-RNA ratio) is applicable to real world consortia, we must first sample microbial communities across transition events and employ quantification techniques that are absolute.

Meta-omics techniques, such as metagenomics (MG)^2,3^, metatranscriptomics (MT)^4^ and metaproteomics (MP)^5^ are routinely used to access prokaryotes in the natural world, where they are part of communities that are frequently dominated by as-yet uncultivated populations^6^. The quantities retrieved from the meta-omics are usually expressed in relative terms, which makes comparison between samples and between omic layers inaccurate^7,8^. Moreover, within dynamic data measurements, such as the MT or MP, the notion of steady state becomes relevant as it is extremely rare that parameters (e.g. bacterial growth rate and nutrient availability) are stable over time^8^.

Here, we present an absolute temporal multi-omic analysis of a minimalistic biogas-producing consortium (SEM1b), which was resolved at the strain level and augmented with two strain isolates^9^. We combined both a RNA-spike-in for MT^10,11^ and the *total protein approach* for MP^12^ for the absolute quantification of high-throughput data. We not only demonstrate that temporal SEM1b samples were comparable within the same omic layer, but also between the MT and MP. Indeed, the protein-to-RNA ratio per sample of the bacterial populations matched previous calculations for the existing example from axenically cultured *E.coli^1^*. For the first time, we present protein-to-RNA ratios for the Archaeal kingdom (*Methanothermobacter thermoautotrophicus*), which are similar to those reported for the Eukarya, and support crystallography and homology studies that suggest the translation system of archaea more closely resembles eukaryotes^13^. Our approach enabled us to explore the linearity of the protein-to-RNA ratio and if it is influenced by changes in community state and/or specific population lifestyle. Finally, we estimated the translation and protein degradation rates, showing that a downregulation of the former marks main lifestyle changes (e.g. motility/chemotaxis and metabolism) during the community development.

## Results and Discussion

### Taxonomic and functional resolution of the omics

In order to characterize RNA/protein dynamics in a microbiome setting, we first needed to molecularly reconstruct our test community over time. Previous analysis of the simplistic SEM1b community genomically reconstructed and resolved 11 metagenome assembled genomes (MAGs) as well as two isolate genomes^9^, covering the taxonomic and functional niches that are required to convert cellulosic material to methane/CO_2_ in an anaerobic biogas reactor^14^. Taxonomic analysis of SEM1b inferred population-level affiliations to *Rumini(Clostridium) thermocellum* (RCLO1), *Clostridium sp.* (CLOS1), *Coprothermobacter proteolyticus* (COPR1, BWF2A, SW3C), *Tepidanaerobacter* (TEPI1-2), *Synergistales* (SYNG1-2), *Tissierellales* (TISS1), and *Methanothermobacter* (METH1)^9^. Herein we estimated that the total genomic potential of SEM1b includes 39144 Open Reading Frames (ORFs) (Supplementary Table 1). Since ORFs with very high sequence similarity may produce RNAs and proteins that are indistinguishable in MT and MP data, we instead gathered all ORFs into ORF-groups (ORFGs), where a singleton ORFG is defined as a group with a single ORF, and thus a single gene. Using this approach, our MT and MP data identified 12552 (96% singleton) and 3235 (78% singletons) highly transcribed and translated ORFGs, respectively. The discrepancy between the singleton percentages was as expected, due to the fact that variations in the DNA/RNA sequences are expected to be greater than in the protein since different codons can code for the same amino acid (codon degeneracy). Degeneracy implies that the chance to distinguish between homologous genes using MT is greater than using MP. Previous MG analyses using assembly algorithms has shown that problematic genomic regions in a given environmental contig can harbor variants from multiple, closely-related strains, which can be further linked to normal strain-level variability within a population and speciation^15–17^. Within SEM1b, the ORFGs that contained multiple homologous ORFs predominantly originated from several strains of a single species. For example, in the MT, 444 non-singleton ORFGs (88% of the total) contained ORFs from different strains of the same species, whilst this was the case for 294 ORFGs (32%) in the MP.

All ORFs were annotated using Kegg Ontology (KO), and at least one term was found for 19070 (49%) representatives from our complete dataset (Supplementary Table 2). The predominant ORF annotations included *Membrane transport*, *Carbohydrate metabolism*, *Translation*, *Amino acid metabolism* and *Replication and repair* (Supplementary Fig. 1). As expected, these functional categories were also among the top five most abundant for the MT, and top six in MP (plus *Energy metabolism*), although in a different order. The *Membrane transport* category is extremely poorly represented in the MP (2% of the terms), which is likely explained by well-known technical issues that limit the extraction of transmembrane proteins^18^. The most abundant annotation categories mentioned above are all in line with the community function of cellulose degradation. The abundance ranking of the KO categories changes slightly from MG to MT (Kendall tau: 0.77, p<10^−8^) and from MT to MP (tau 0.74, p<10^−6^) whilst moderately from MG to MP (tau 0.68, p<10^−5^), which means that the functional potential observed in the genomes is more preserved in the diversity of produced transcripts than the one of proteins and thus hints to post-transcriptional regulation playing an important role in addition to transcriptional regulation in prokaryotes.

### Absolute quantification extends expectation from *E.coli* RNA/protein dynamics and positions Archaea alongside the Eukarya

To determine whether or not microbial RNA/protein dynamics vary between ecological status (isolate vs community), metabolic states and/or taxonomic phylogeny, we quantified and resolved the numbers of transcript and protein molecules per sample in our SEM1b community, which averaged 3.8×10^12^(sd 3.0×10^12^) and 2.2×10^15^(sd 9.5×10^14^), respectively (Supplementary Tables 3-4). Microbial cell volume and its transcriptome size has been shown to change in yeast according to cell status (proliferation vs. quiescence), whilst the proteome is merely reshaped in its composition^19^. In our case, the number of total transcripts per SEM1b sample increased more than three-fold during the first 15 hours (from ~1.2×10^12^ in t1 to ~4.0×10^13^ in t4) in the SEM1b consortium’s life cycle and then decreased sharply, whereas the number of proteins per sample reached a plateau after 18 hours post-inoculation at ~2.7×10^15^ molecules. SEM1b approximated the exponential growth phase in t3 (18 hours), therefore we used the protein-to-RNA ratio from this time point for comparison against previously reported axenic estimates^1,20–23^. The replicate-averaged protein-to-RNA ratio for the bacteria in SEM1b ranges from ~10^2^ to 10^4^ (median = 949, Fig. 1a), which fits the estimated range reported for *E.coli^1^*. This means that for every RNA molecule one can expect from 100 to 10000 protein molecules with a value of 949 being the most likely. Our results showed a taxon-specific variation in the protein-to-RNA ratio within bacteria (Fig. 1a). Indeed the median ratios for the bacteria in SEM1b at 18h ranged from 658 in CLOS1 to 1137 in RCLO1. Moreover, we report, for the first time, the median protein-to-RNA ratio for an Archaeal organism: METH1 (*M. thermoautotrophicus*) as being 12035 protein molecules per detected RNA (Fig. 1a). The reported values for Eukaryotes are 4200-5600 for yeast^20,21^ and 2800-9800 for *Homo sapiens^22,23^*; therefore, we find that Archaeal translation dynamics are closer to that observed within the Eukaryotic kingdom than that of bacteria.

**Figure 1.**
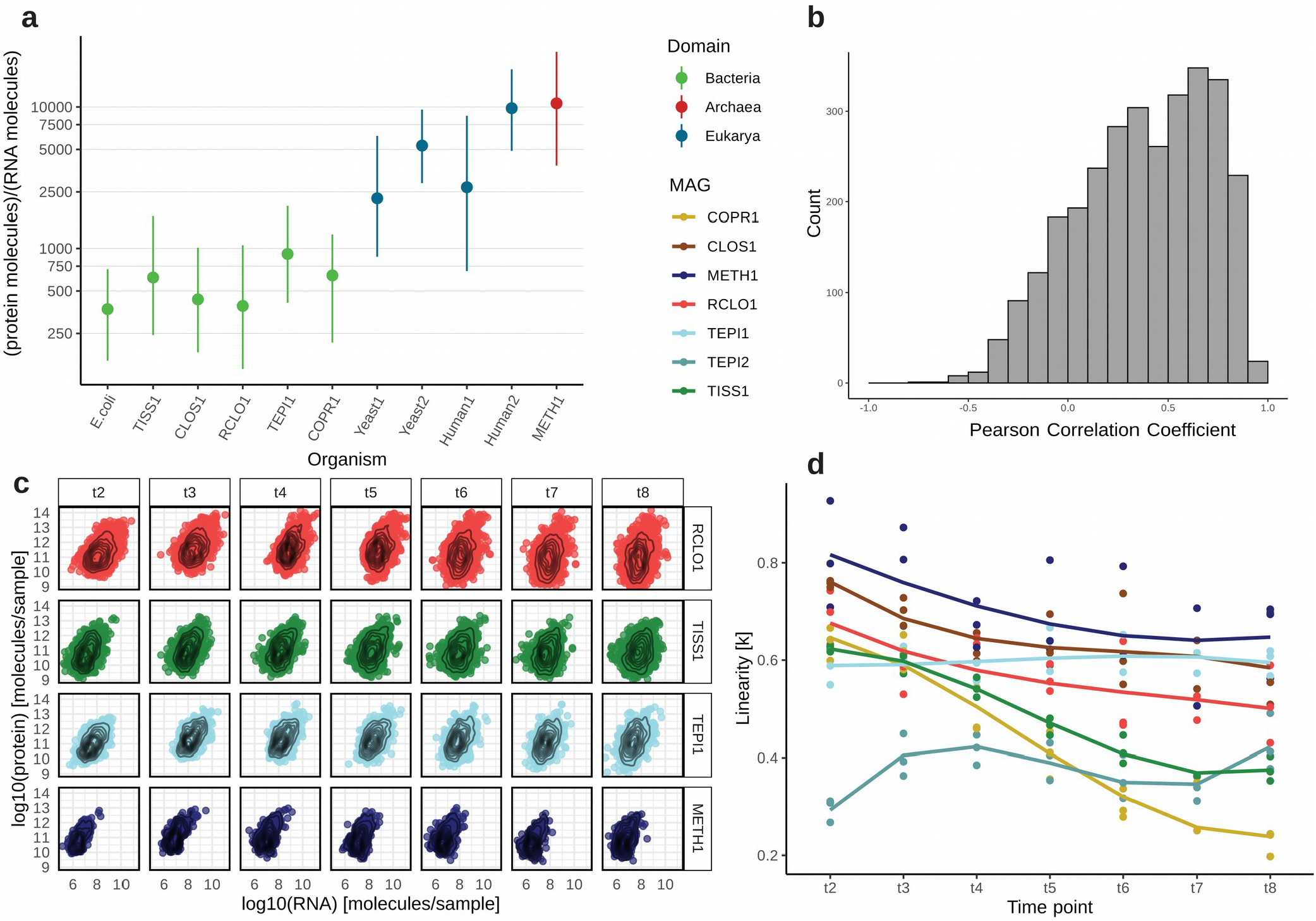
Protein-to-RNA ratio distributions of as-yet uncultured bacterial and archaeal populations within a microbial community. **a.** Comparison of protein-to-RNA ratio distributions of selected MAGs reconstructed from the SEM1b community as well as those previously reported in the literature. The dots represent the median values and the bars span from the first to the third quartiles. The protein-to-RNA ratios for *E.coli* was retrieved from Taniguchi et al.^1^, Yeast1 from Ghaemmaghami et al.^20^, Yeast2 from Lu et al.^21^, Human1 from Schwanhausser et al.^22^ and Human2 from Li et al.^23^. **b.** The distribution of the Pearson Correlation Coefficients (PCC) between transcripts and their corresponding proteins computed across the time points. With a median PCC of 0.41, the change in the amount of a given transcript over time seemingly does not translate into a change in the amount of the corresponding protein. **c.** Per-time-point scatterplots of the absolute protein and transcript levels for ORFs that produced both detectable transcript and protein in SEM1b datasets. For simplicity, only four representative MAGs are shown, with all MAGs depicted in Supplementary Fig. 2. **d.** The plot shows how the linearity parameter *k* between RNA and protein changes over time for the different MAGs. The linearity represents how a change in RNA level is reflected in a change in the corresponding protein level. The parameter ranges from 0 to 1, and increasingly smaller values translate in fewer proteins being expected for the same level of RNAs. The populations CLOS1, METH1 and TEPI1 are converging towards the same values, while RCLO1 has a parallel trend. Hinting to the existence, and the reaching of an equilibrium among them.

A bacterial cell is considered to be in steady state during the log phase of its growth cycle ^8,24,25^, specifically when the changes in proteome size are believed to be mainly dictated by a change in the transcriptome^26^. In contrary to these assumptions, comparisons of RNA and protein levels between individual cells of *E. coli* grown at steady state have not been shown to correlate, however patterns do emerge when all cells are collectively considered at the population level^1^. In SEM1b, we wanted to see if correlations between RNA and protein levels exist in a larger microbial community, and if they are affected by changes in time and life stages. We calculated gene-wise Pearson Correlation Coefficients (PCCs) of protein and transcripts over time for all SEM1b populations and showed that the PCC value varied greatly (Fig. 1b) with a median of 0.41, suggesting that no direct correlations between RNA and proteins levels exist at any stage in a microbiome and that it is nearly impossible to predict the level of the given protein based on the level of the corresponding transcript.

Looking at relationships between proteome and transcriptome for individual populations within SEM1b (examples form four populations in Fig. 1c) was observed to follow a more predicable relationship, which can be described by the monomial function:

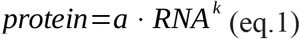

The formula for log10-transformed RNA and protein levels takes the form of a linear model (see methods) that was fitted to protein and RNA distributions per time point from MAGs with the highest quality (RCLO1, CLOS1, COPR1, TISS1, TEPI1, TEPI2 and METH1) (Fig. 1d). The linearity parameter k can be interpreted as the rate of which a change in RNA level is reflected in a change in the corresponding protein level. For example, if k=1, a doubling in RNA level means a doubling in protein level, whereas if k=0.5 a doubling in RNA level means a ~40% increase in protein level. Ranging from 0 to 1, it implies that, in the “perfect” condition where k=1, the number of proteins is linked to the number of RNAs by the scalar constant *a*, whilst if k approaches 0, there will be much lower expected protein levels for the same number of RNAs. With the exception of TEPI2, the linearity (*k*) between protein and RNA levels was observed to start at values between 0.6 and 0.8 at 13 hours (t2) (Fig. 1d). The evolution of the MAGs’ *k* values over time is then divided in three groups: one which is losing linearity rapidly (TISS1 and COPR1); one which is slowly declining (RCLO1, CLOS1 and METH1) and one which is staying constant if not increasing (TEPI1 and TEPI2) (Fig. 1d). Notably CLOS1, METH1 and TEPI1 are converging towards the same linearity values, while RCLO1 has a parallel trend to them. If these trends can be used to retro-fit the steady state definition, we can hypothesize that these four populations possess a metabolic equilibrium and that this equilibrium is approximately reached within the 10 hour window between 33h and 43h (t6 and t7 respectively, Fig. 1d).

### Interpretation of functional specialization in the light of RNA-protein dynamics

Using multi-omic data and the above described RNA-protein dynamics, we were able to visualize that at least four populations within SEM1b converge upon a dominant metabolic state that we speculate to strongly shape the overall SEM1b community phenotype and suggest a functional co-dependence between the individual populations. To determine if this was the case, we annotated the genes and metabolic pathways for SEM1b MAGs (Fig. 2) and reconstructed their temporal expression patterns (Fig. 3). The SEM1b consortium is able to convert cellulose (and hemicellulose) to methane via the combined metabolism of its seven major constituent populations (Fig. 3a). Based on previous analysis that showed that RCLO1 is closely related to *R. thermocellum^9^*, we predict that it senses^27^ its growth substrate (cellulose) and moves towards it (Fig. 3d). RCLOS1 then transcribes, translates and secretes the components of the cellulosome, such as scaffoldins, dockerins and carbohydrate-active enzymes (CAZymes)^28^, which assemble into a dynamic multi-proteins complex that degrades the substrate to smaller carbohydrates. Via the MG, we predicted that non-cellulosomal CAZymes were also employed by the *Clostridium*-affiliated CLOS1, which acted upon the hemicellulose fraction (mainly xylan) trapped in the spruce cellulose, which was supported by observed release of its main monomer xylose (Fig. 3a). Sugars generated via the actions of RCLO1 and CLOS1 are subsequently consumed by RCLO1, CLOS1 and *Coprothermobacter*-affiliated populations (COPR1, BWF2A and SW3C), which were all observed to express sugar transporters (Fig. 2). Notably CLOS1 has the most diversified transporters, making it a flexible consumer, and for the most part demonstrated highest levels of hydrolytic and fermentative gene expression after RCLO1, which again is likely tied to xylose release at later stages of the SEM1b lifecycle (Fig. 3a). However, some of the transporters, such as the one for oligogalacturonide, raffinose/stachyose/melibiose and rhamnose, were not expressed, likely due to the absence of their substrates in the largely cellulose and xylan dominated spruce wood used in this study. CLOS1 was also the only population to possess the aldouronate transporter with 20 copies of gene lplA, 20 of lplB and 16 of lplC (20/20/16) and expressing 0.4/0.7/3.8×10^10^ and 92.8/3.5/7.0/×10^11^ combined median transcripts and proteins per sample; making it one of the few transporters detectable at the protein level. Similarly, the *C. proteolyticus* strains (BWF2A and SW3C) possess and express unique sugar transporters, likely gaining access to an undisputed pool of arabinogalactan or maltooligosaccharide. The transporter for pentamers ribose/xylose were the most common and possessed by RCLO1, *C. proteolyticus* populations and *Tepidanaerobacter* populations (TEPI1 and TEPI2). Notably from Fig. 2, it is clear that the proteins from the transporters are almost never found in the samples, even if the respective RNAs are abundant. This is likely due to the difficulties in extracting transmembrane proteins^18^.

**Figure 2.**
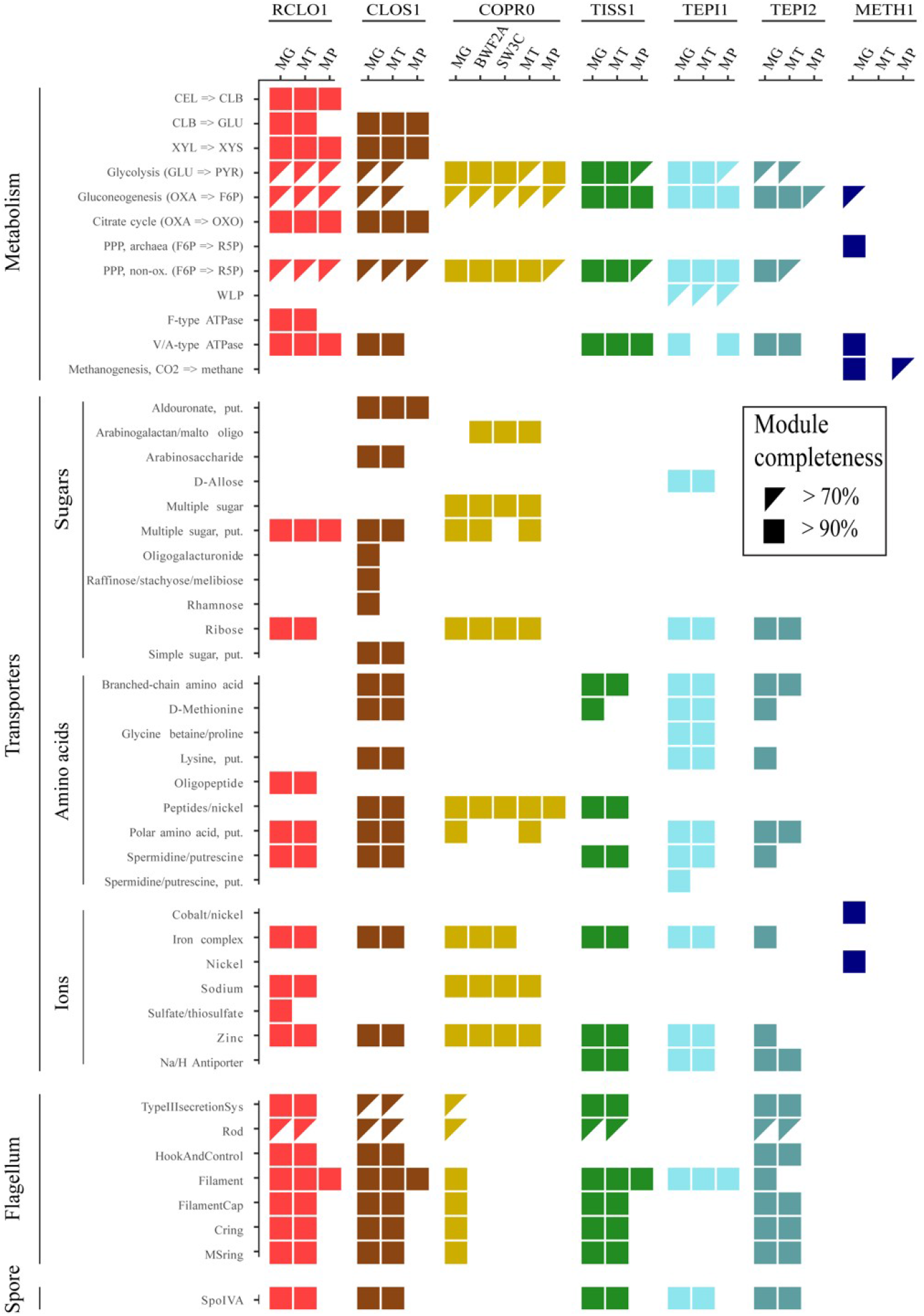
Overview of the genetic potential and expressed modules in the seven populations of SEM1b. Module completeness denotes the level of detected RNA and proteins mapped to major genes/metabolic pathways that are critical to the SEM1b lifecycle. Only MAGs with the highest quality reconstruction (RCLO1, CLOS1, COPR1, TISS1, TEPI1, TEPI2 and METH1) are included as well as two isolated and genome-sequenced *Coprothermobacter* strains, for which the transcriptome and the proteome were considered as the species level.

**Figure 3.**
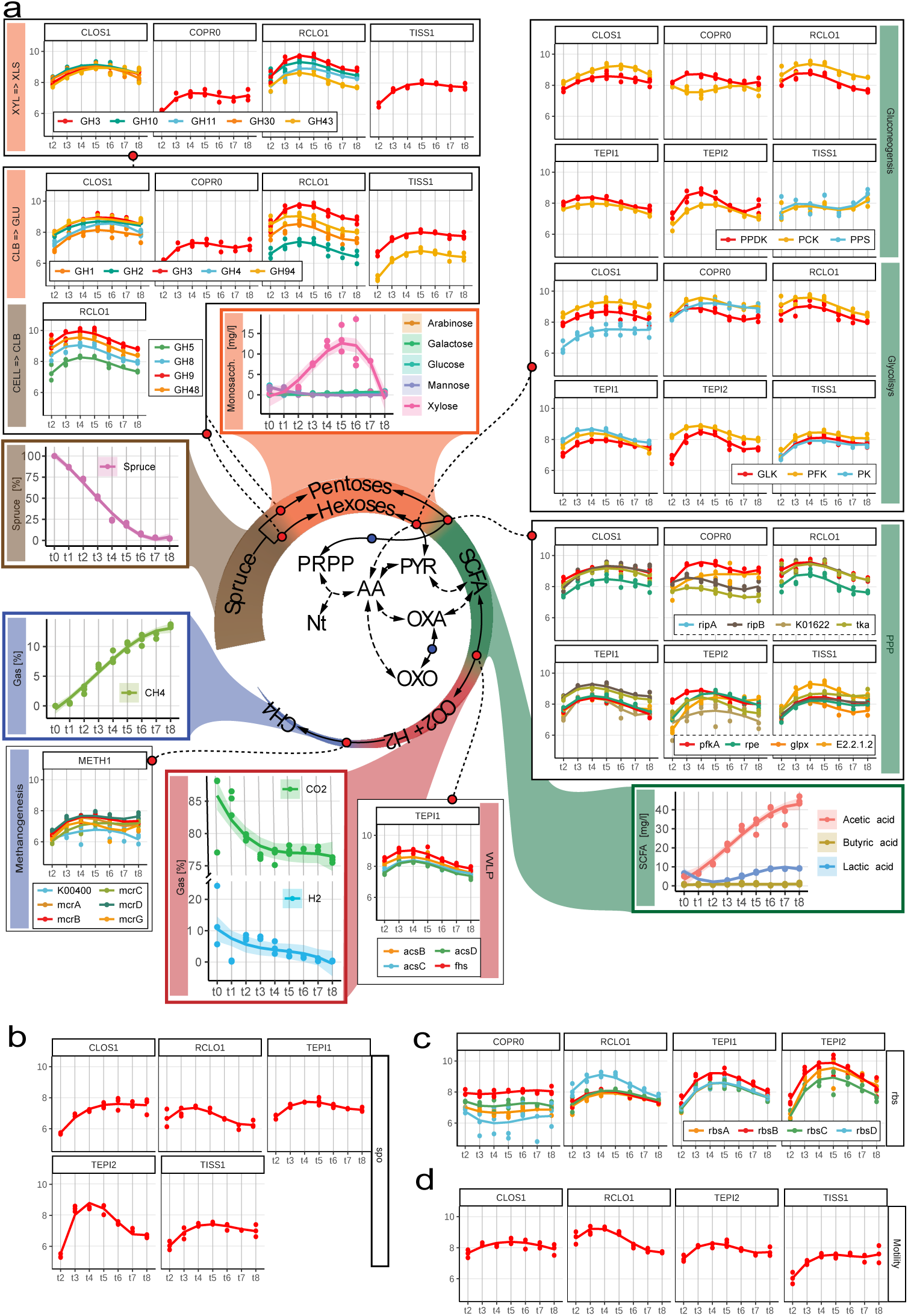
Schematic representation of the temporal and co-dependent metabolism of SEM1b that converts spruce-derived cellulose to methane. **a.** Within SEM1b, four major metabolic stages are required: Spruce → Hexoses/Pentoses, Hexoses/Pentoses → SCFAs, SCFAs → CO_2_+H_2_ and CO_2_+H_2_ → methane). Metabolites (spruce, sugars, SCFAs, CO_2_+H_2_ and methane) involved in these processes were measured and the temporal analysis of the metabolic pathways involved in their interconversion is depicted for the major SEM1b populations. Other metabolites (for which abbreviations are: Nt=Nucleotides, PRPP=Phosphoribosyl pyrophosphate, AA=Amino acids, PYR=Pyruvate, OXA=Oxalacetate and OXO=Oxo-glutarate) are shown to highlight the essential metabolism of the microbes. In the central metabolic network the metabolites are linked by solid arrows if the interconversion requires one step or the link between them is addressed more in detail (blue dot if in Fig. 2, red dot if in a pathway plot herein). Metabolic pathways are quantified via marker genes (selection in methods section) in the scale of log10-transformed transcript molecules per sample whilst the solid lines in the plots represent the qubic fitting of the data points. More metabolites’ abbreviations are CELL=Cellulose, CLB=Cellbiose, GLU=Glucose, XYL=Xylan, XLB=Xylobiose and pathways’ abbreviations are WLP=”Wood-Ljungdahl Patway”, PPP=”Penthose Phosphate Pathway”. **b.** Sporulation is common to all Gram positive bacteria of the community and it is quantified with the marker *spoIVA*. Notably TEPI2 is investing greatly in spore formation until 28h after the inoculum (t4). **c.**The genes for the Ribose and xylose transporter (*rbs*) are expressed in four populations. Notably TEPI2 produces more rbs transcripts than the closely related MAG TEPI1; indeed, the first has been predicted to be a mere fermenter whilst the latter bases its metabolism on the WLP pathway (Fig. 3a). **d.** Microbial motility is represented by the marker gene flgD. RCLO1 is the most active bacterium, producing less and less flagella over time after t4. It starts ahead of the others at t2, presumably finishing the colonization of the substrate (Spruce-derived cellulose).

The process of degrading cellulose and simple saccharides via hydrolysis and fermentation ultimately results in the production of short chain fatty acids (SCFAs) such as proprionate, butyrate and acetate, which are subsequently metabolized by the SCFA-oxidizing populations in SEM1b (TISS1, TEPI1, TEPI2) (Fig. 3a). The only metabolically-active SCFA-oxidizing population in SEM1b was predicted to be TEPI1, as it demonstrated good linearity between protein and RNA levels that increased over time (Fig. 1d) and harbored a complete Wood-Ljungdahl Pathway (WLP) that was detectable in both MT and MP (Fig. 2). It has been shown that oxidizers can improve oxidization of SCFAs (up to double speed) when superior NADPH and ATP generators (e.g. glucose) are consumed in small amounts to complement the stoichiometry through the Pentose Phosphate Pathway (PPP) without triggering the shift of the entire cells metabolism toward another substrate^29^. In this context, it is interesting to note that TEPI1 was the only MAG that encoded and expressed a hexose (allose) transporter (Fig. 2). Aldohexoses (such as D-allose, D-glucose, D-mannose, etc.) are imported and transformed into fructose-6P in two reactions (both expressed in TEPI1), which can then be fed into both the PPP or the Glycolysis pathways. Xylose, is a product of the degradation of hemicellulose present in our system (Fig. 3a) and can be converted to ribulose-5P and fed to the PPP in three reactions. This data, in combination with a highly expressed and detectable WLP over time (Fig. 3a), points to the establishment of TEPI1 as the only syntrophic acetate-oxidizer (SAO) in the SEM1b consortium. We speculate that TEPI1’s SAO-metabolism is helped by the other SEM1b populations that generate acetate as a fermentation end-product and the supplement of the sugars released by the cellulosomal complex such as glucose and xylose. Interestingly the closely related MAG TEPI2 was observed to lack the WLP and to express ~10 times more transcripts for the ribose/xylose transporter than TEPI1; relegating it to the role of mere sugar degrader, and probably scavenger in the community.

While TISS1 seems mostly to phase out of the community and lose linearity in its protein to transcript relationship (Fig. 1d), TEPI2 implements an exit strategy in the form of sporulation. All the gram-positive populations from the SEM1b consortium (RCLO1, CLOS1, TISS1, TEPI1 and TEPI2) were able to produce spores and express the spore marker *spoIV*, an ATPase associated to the surface of the neospore that promotes the assembly of the coating and is common to all the spore forming bacteria^30^ (Fig. 3b). TEPI2 however increased the level of transcripts for spoIV by 1000 times within the 13h and the 18h time points, reaching the maximum at 23h, and having a production 10 times higher than the phylogenetically related TEPI1. All SEM1b populations, except the *C. proteolyticus* isolates and TEPI1, have the genetic potential for flagellar synthesis but the respective transcripts were only observed for RCLO1, CLOS1, TISS1 and TEPI2. The filament protein of RCLO1 is by far the most abundant protein in the samples with an average of 2.8×10^13^ molecules per sample, which matches the idea of RCLO1 investing in motility to reach the cellulose fibers and starting with the highest level of marker flgD in the community (Fig. 3d).

In microbial ecosystems, acetate oxidization is a syntrophic process, whereby end-products of the WLP CO_2_ and H_2_ /formate are co-metabolized by hydrogenotrophs such as Archaeal methanogens. The methanogenesis pathway encoded in METH1 is the largest pathway according to the number of genes involved (n=112) in SEM1b, while we also observed transporters for nickel, the metallic ion used by the nickel-containing methyl-coenzyme M reductase (the central enzyme in methanogenesis^31^). Methanogenesis also use electrochemical gradients generated by Na^+^ and H^+^ ions to drive energy production and recharge the electron donor groups (ferrodoxin, F420), similar to SAO bacteria. Peculiarly, the populations TEPI1, TEPI2 and TISS1 were the only ones found to encode and express the Na^+^/H^+^ antiporter *nha* (Fig. 2) pointing to an important role of these ions in the greater SEM1b consortium. The WLP is associated with the transition between NADH/NAD^+^, and translocates Na^+^ to create a gradient, which is used by the type-V ATPase to synthesize ATP^32^. Indeed the NAD^+^-Fd_Red_-dependent Na^+^ translocation system *rnf* is expressed in both the fermenting and SAO bacteria of SEM1b, while type-V ATPase, which produce energy by exploiting the Na^+^/H^+^ gradient, were detected by all the populations aside from METH1 and *C. proteolyticus* (Fig. 2). Moreover, the TEPI1 MAG expresses the HND NADP-reducing hydrogenases complex, which turns hydrogen ions into H_2_ using NADPH. The molecular hydrogen can then permeate the membrane and be used by the syntrophic partner METH1 to generate methane (Fig. 3a).

### Translation control drives changes in cell status and source utilization

In addition to RNA/proteins ratio assessments, our collection of absolute multi-omic data allowed us to explore the crucial aspect of protein-level regulation, which is poorly understood in microbiomes. The control of protein levels in bacteria is believed to occur predominately via transcription control, “control by dilution”^33^ (dispersal of proteins via subsequent cell divisions) and rarely by protein degradation^34^. Similar to transcription control, translation can also be controlled by a dynamic pool of translational factors, such as initiation, elongation and ribosome components^35^. The processes targeted by these systems require a rapid change in the number of proteins in the cell that cannot wait for a change in RNA levels or a dilution effect. The absolute quantification of transcripts in SEM1b and proteins was used to estimate the translation and protein degradation rates using PECA-R^36^ (Supplementary Table 5). The analysis found 305 significant changes in translation rate, accounting for 302 ORFs. Of the rate changes’, 94% were downregulated and 71% of the ORF were functionally annotated. RCLO1 has 28 downregulated ORFs between 13h and 18h (t2-t3), mostly from complexes involved in chemotaxis (*cheY*, *cheW*, *mcp*), flagellum assembly (*flgG*, *flgK*, *fliD*) and shape determination (*mreB*). In the following five hours, several systems concerning carbon fixation are affected, such as phosphoglicerate kinase (PGK), triosephosphate isomerase (TPI), phosphate acetyltransferase (EC 2.3.1.8), isocitrate dehydrogenase (IDH1) and *pyruvate* orthophosphate dikinase (PPDK). In the next five hours it downregulates the translation of the cell division protein zapA as well. The reduction protein production for chemotaxis, mobility and then cell division matches the idea that within 13h of the inoculation, RCLO1 sensed, reached and colonized the cellulose fibers. Contextually the release of medium length carbohydrates enables RCLO1 to engage in the more energetically favorable fermentation metabolism. TISS1 has a decrease in translation rates of ORFs related to metabolic processes between 13h and 18h, mostly involving cofactors (*fhs*, *folC*, *folD*, *lplA*, *metH*, *pdu0* and *nadE*) and amino acids (*aorQ*, *hutI*, LDH, *metH*, *mtaD* and *pip*). TEPI1 down-regulated 60 ORFs, accounting for part of its carbohydrate metabolism (e.g. PGK, TPI), the amino acid transporters and the NADH dehydrogenase complex (HND). TEPI2 has 19 ORFs subject to downregulation in the 13h-18h interval, such as Pyruvate ferrodoxin odidoreductase (PFOR), GK, fructose-bisphosphate aldolase (FBA), tansaldolase EC 2.2.1.2 and the ribose/xylose transporter subunit *rbsB*. In the last interval (33h-38h), RCLO1 upregulated the translation of 10 ORFs, among which the flagellar flbD and shape determination mreB; seemingly starting to restore the functions downregulated in the 13h-18h interval.

## Conclusions

We present the reconstruction of a microbiome from a model environment and quantified the number of RNAs and proteins over time in absolute terms. This approach enabled us to assess and report, for the first time, the protein-to-RNA ratio of multiple microbial populations simultaneously, which individually engage in distinct, yet integrative metabolic pathways that ultimately cumulate into the community’s principal phenotype of converting cellulose to methane. We extended the results from Taniguchi et al.^1^, showing that our populations had a varying protein-to-RNA ratio in the predicted interval of 10^2^-10^4^ while presenting for the first time the same quantity for an Archaeal population (METH1): 10^3^-10^5^, which resembled the previously measured values for Eukaryotes^20–23^. The greater ecological significance of the seeming Archaeal capacity to generate higher protein levels at a lower “RNA-cost” is of interest, as many Archaeal populations in mixed-kingdom microbiomes are known to exert strong functional influence, despite their cell concentrations being orders of magnitude lower than their bacterial counterparts (i.e. the rumen microbiome^37^).

Moreover, we assessed the linearity between transcriptome and proteome for each population over time (Eq.1), finding that three major populations of the community, a fermenter (CLOS1), a SAO (TEPI1) and a methanogen (METH1), were converging on the same values in parallel with the primary cellulose degrader (RCLO1) (Fig. 1d). The highlight of their seemingly intertwined protein/RNA dynamics matches with their metabolic complementarity, starting from RCLO1 degrading cellulose to sugars and SCFAs, CLOS1 fermenting sugars to SCFA, TEPI1 oxidizing SCFAs to H_2_ and METH1 converting CO_2_ and H_2_ to methane. Closer examination revealed even more intricate relationships involving Na^+^ and H^+^ ions as well as secondary sugars (i.e. xylose) reiterating that each population needs the metabolic activity and subsequent byproducts of the previous one to provide a supply of growing metabolites (Fig. 3a). Moreover, the estimation of translation and protein degradation rates pointed at a translational negative control for several ORFs involved in chemotaxis/motility and central metabolism, marking important changes in the community status. In conclusion, our data highlights that simple modifications to multi-omics toolkits can reveal much deeper functional-related trends and integrative co-dependent metabolisms that drive the overall phenotype of microbial communities, with potential to be expanded to more-complex and less-characterized microbial ecosystems.

### Data availability

All sequencing reads have been deposited in the sequence read archive (SRP134228), with specific numbers listed in Supplementary Table 6 in Kunath et al.^9^. All microbial genomes are publicly available on JGI under the analysis project numbers listed in Supplementary Table 6 in Kunath et al.^9^. The mass spectrometry proteomics data have been deposited to the ProteomeXchange Consortium via the PRIDE^38^ partner repository with the dataset identifier PXD016242. The code used to perform the computational analysis is available at: https://github.com/fdelogu/SEM1b-Multiomics.

## Supporting information

Summary of supplementary figures and tables

## Acknowledgements

We are grateful for support from The Research Council of Norway (FRIPRO program, PBP: 250479), as well as the European Research Commission Starting Grant Fellowship (awarded to PBP; 336355 - MicroDE). The sequencing service was provided by the Norwegian Sequencing Centre (www.sequencing.uio.no), a national technology platform hosted by the University of Oslo and supported by the “Functional Genomics” and “Infrastructure” programs of the Research Council of Norway and the Southeastern Regional Health Authorities.

## Author contributions

F.D., T.R.H. and P.B.P. conceived the study, performed the primary analysis of the data and wrote the paper (with input from all authors). B.J.K., and M.Ø.A. generated the data and contributed to the data analyses.

## Competing interests

The authors declare there are no competing financial interests in relation to the work described.

## Materials and Methods

### Multiomics data acquisition

#### Background

The full experimental setup and the methods concerning the retrieval of biological samples and data preprocessing were performed during a previous study^9^ and can be summarized as follows: a microbial consortium called SEM1b was obtained from a biogas reactor using serial dilution and enrichment methods on spruce cellulose. A metagenomic analysis was initially performed on the SEM1b community using two different generations that had consistent population structure, and was used as a supporting database for a subsequent SEM1b time series experiment. The time series analyses consisted of metabolomics, metaproteomics and metatranscriptomics over nine time points (at t0, 8, 13, 18, 23, 28, 33, 38 and 43 hours) in triplicate (A, B and C), spanning the consortium life-cycle.

#### Metagenomics

For generation of metagenomic data, 6ml samples of SEM1b culture were taken and cells were pelleted prior to storage at −20°C. Non-invasive DNA extraction methods were used to extract high molecular weight DNA as previously described in Kunath et al.^39^. DNA samples were prepared with the TrueSeq DNA PCR-free preparation, and sequenced with paired-ends (2×125bp) on one lane of an Illumina HiSeq3000 platform (Illumina Inc) at the Norwegian Sequencing Center (NSC, Oslo, Norway). Metagenomic analyses comprising quality trimming and filtering, reads assembly, binning and annotations were performed as previously described^9^. Resulting annotated open reading frames (ORFs) were retrieved and used as a reference database for the metatranscriptomic and metaproteomic analysis.

#### Metatranscriptomics

mRNA extraction was performed in triplicate on time points t2 to t8, using previously described methods^11^. The extraction of the mRNA included the addition of an in vitro transcribed RNA as an internal standard to estimate the number of transcripts in the natural sample compared with the number of transcripts sequenced. For further normalization, total RNA was extracted using enzymatic lysis and mechanical disruption of the cells and purified with the RNeasy mini kit (Protocol 2, Qiagen, USA). The RNA standard (25ng) was added at the beginning of the extraction in every sample. After purification, residual DNA, free nucleotides and small RNAs were removed. Samples were treated to enrich for mRNAs and then amplified before being sent for sequencing at the Norwegian Sequencing Center (NSC, Oslo, Norway). Samples were subjected to the TruSeq stranded RNA sample preparation, which included the production of a cDNA library, and sequenced with paired-end technology (2×125bp) on one lane of a HiSeq 3000 system.

The resulting sequences were filtered and rRNA and tRNA reads were removed as performed in Kunath et al.^9^. The reads mapping on the internal standard pGEM-3Z were extracted using SortMeRNA^40^ v2.1b and their counts used as *I*_*R*_ in the “Functional omics absolute quantification” section of the material and Methods, whilst the not mapping reads (the transcriptome in the sample) were used as ∑*T*_*R*_. The retained reads were mapped against the predicted genes dataset using Kallisto pseudo-pseudobam^41^ and the mapping files were produced with bam2hits. Transcripts were quantified with mmseq^42^ and collapsed using mmcollapse^43^.

#### Metaproteomics

Proteins were extracted from t1 to t8 in triplicate following a previously described method^44^ with a few modifications. Briefly, 30ml of cultures containing cells and substrate were centrifuged at 500x g for 5 minutes to pellet the substrate. The supernatant was centrifuged at 9000 x g for 15 minutes to collect the cells. Cell lysis was performed by resuspending the cells in 1ml lysis buffer (50 mM Tris-HCl, 0.1% (v/v) Triton X-100, 200 mM NaCl, 1 mM DTT, 2mM EDTA) and keeping them on ice for 30 minutes. Cells were disrupted in 3 × 60 seconds cycles using a FastPrep24 (MP Biomedicals, USA). Debris were removed by centrifugation at 16000 x g for 15 minutes. The supernatants containing the proteins were kept at −20°C until further processing. Extracted proteins were quantified using the Bradford’s method. 50μg of each sample were denatured using SDS sample buffer and loaded on an Any-kD Mini-PROTEAN gel (Bio-Rad Laboratories, USA) and separated by SDS-PAGE for 20 minutes at 270V. Each gel lane was cut into 16 slices and the reduction, alkylation and tryptic digestion of the proteins into peptides were performed in-gel. The tryptic peptides were extracted from the gel and desalted prior to mass spectrometry analysis. Peptides were analyzed using a nanoLC-MS/MS system connected to a Q-Exactive hybrid quadrupole-orbitrap mass spectrometer (Thermo Scientific, Germany) equipped with a nano-electrospray ion source. The Q-Exactive mass spectrometer was operated in data-dependent mode and the 10 most intense peptide precursors ions were selected for fragmentation and MS/MS acquisition. The selected precursor ions were then excluded for repeated fragmentation for 20 seconds. The resolution was set to R=70,000 and R=35,000 for MS and MS/MS, respectively.

A total of 384 raw MS files (8 samples x 3 biological replicates x 16 fractions) were analyzed using MaxQuant^45^ version 1.4.1.2 and proteins were identified and quantified using the MaxLFQ algorithm^46^. The data was searched against the generated MG dataset from Kunath et al.^9^ supplemented with common contaminants such as human keratin and bovine serum albumin. In addition, reversed sequences of all protein entries were concatenated to the database for estimation of false discovery rates. The tolerance levels for matching to the database was 6 ppm for MS and 20 ppm for MS/MS. Trypsin was used as digestion enzyme, and two missed cleavages were allowed. Carbamidomethylation of cysteine residues was set as a fixed modification and protein N-terminal acetylation, oxidation of methionines, deamidation of asparagines and glutamines and formation of pyro-glutamic acid at N-terminal glutamines were allowed as variable modifications. The ‘match between runs’ feature of MaxQuant^46^ was applied. All identifications were filtered in order to achieve a protein false discovery rate (FDR) of 1%. Quantitative information was retrieved using the LFQ intensities of each proteins.

#### Metabolomics

For monosaccharide detection, 2 ml samples were taken in triplicates, filtered and sterilized with 0.2μm sterile filters and 15 minutes boiling. Soluble sugars were identified and quantified by high-performance anion exchange chromatography (HPAEC) with pulsed amperiometric detection (PAD). For quantification, peaks were compared to linear standard curves generated with known concentrations of selected monosaccharides (glucose, xylose, mannose, arabinose and galactose) in the range of 0.001-0.1 g/L.

For the short chain fatty acids (SCFAs), 1ml was taken in triplicate from each time point, they were centrifuged at 16000 x g for 5 minutes and the supernatants were filtered with 0.2μm sterile filters. 5μL of Sulfuric Acid 72% were added to the filtrates and let at rest for 2 minutes before being centrifuged again at 16000 x g for 5 minutes, transferred in a new tube and stored at −20°C until processing. SCFAs were then analyzed using a Dionex 3000 HPLC as described in Estevez et al.^47^.

### Functional omics absolute quantification

#### Metatranscriptomics

The absolute quantification of transcripts was taken from Mortazavi et al.^10^ using the internal standard from Gifford et al.^11^ as reference to estimate the length of the initial transcriptome. The number of reads produced in a given sample is proportional to the total amount (in Nt) of starting material. With the addition of an internal standard we have the following proportion between the starting material for transcripts (T_Nt_) and the internal standard (I_Nt_) and the reads they produce (T_R_ and I_R_ respectively):

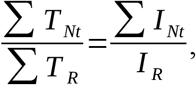

in which the sums are taken over a single sample. The formula can be rearranged as:

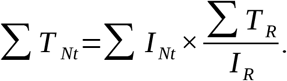

Since we know the number of molecules of internal standard added (I_M_) and its length (I_Nt_), we can substitute them in the equation as:

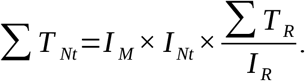

We can now use the estimation of the starting length of the transcriptome and the TPMK transcript measurements in the formula from Mortazavi et al.^10^:

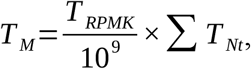

which becomes:

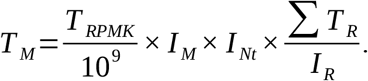

#### Metaproteomics

The “Total protein approach” method from Wiśniewski et al.^12^ relies on the use of the protein mass per sample, the computed Molecular Weight (MW) of the detected proteins to transform the LFQ values into absolute ones. Here we omitted the per-cell quantification since SEM1b is a heterogeneous community and MG measurements were not taken for the time series.

We computed the *Total protein*_*i*_ as:

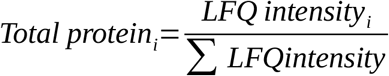

Then the *Protein concentration*_*i*_ was obtained from the previous with:

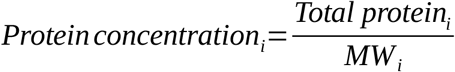

The method was developed on the assumption that the reference proteome is complete and that the total mass of the peptides detected is equal to the total mass of peptides processed by the machine. This is not necessarily valid in a microbiome for which the reference cannot be completely reliable. Thus we computed the fraction of identified mass using the raw MP files with the following formula:

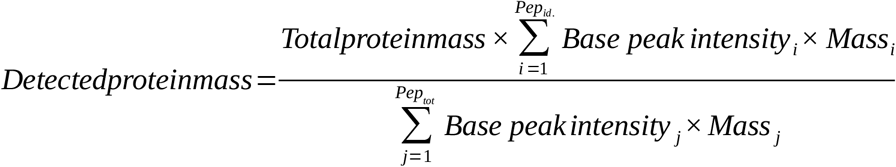

Finally the copy number of proteins per sample was computed using the Avogadro’s Number (N_A_) as:

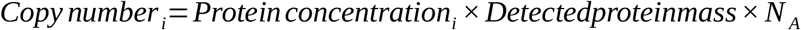

### Multiomics dataset integration

#### Data preprocessing

The MT and MP datasets estimate absolute abundance of ORFGs over time. An expression group is defined in this study as a set of ORFs which cannot be further resolved using the available data. When the analysis required the direct comparison of ORFs (e.g. transcript-protein correlation) only the singleton subset of the ORFGs was considered. The reliability of the expression estimation is linked to the number of unique hits (reads or peptides) available for a given ORF, therefore all the entries with 0 unique hits were filtered out. The datasets were then log10-transformed with a pseudocount equal to one. After expression density plotting, the minimum expression thresholds of 5 and 9 were selected for MT and MP, respectively, and the data was filtered accordingly. Principal component analysis was used to screen the samples and t7C (time point 7, replicate C) was identified as an outlier and removed before downstream analysis.

#### MP/MT linear fit

We took the intersection of ORFs present in the MT and MP layers of the dataset for each of the selected MAGs (COPR1, CLOS1, COPR1, METH1, RCLO1, TEPI1, TEPI2, TISS1), and, for each sample, we performed a regression analysis in R. The values span several orders of magnitudes, thus we decided to fit the monomial functional:

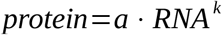

which can be rewritten as:

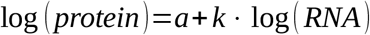

to be more easily fitted as a linear model. The previously log10 transformed protein levels were used as *y* while the log10-transformed RNA was used as *x* in a linear model using the lm function. The slopes of the models were then used to fit a third grade polynomial function to obtain the linearity change profile in Fig. 1d.

#### Functional annotation and module completeness

The KEGG Orthology (KO) numbers were assigned to the ORFs as a part of the annotation pipeline from IMG^48^. The ORF-wise annotation was then translated into the MT/MP-ORFGs assigning to each ORFG a non-redundant set of all the terms assigned to all the ORFs in the group. We used the KO numbers to estimate the KEGG module completeness using the R package MetaQy^49^ v.1.1.0. The Glycosyl Hydrolases annotation was retrieved from Kunath et al.^9^.

#### Metabolic marker genes selection

The metabolic marker genes for Fig. 2 were selected with the following criterion. Glycolysis/Gluconeogenesis: enzyme with irreversible reactions. PPP: genes involved in the main interconversion loop between Ribose-5 Phosphate and Fructose-6 Phosphate. WLP: marker genes from Can 2014. Methanogenesis: markers genes from Scheller 2010. The Glycosyl Hydrolases were manually curated to assemble a set able to perform the substrate conversion.

#### PECA analysis

We ran PECA-R^36^ to estimate translation and protein degradation rates using the absolute quantification tables for transcripts and proteins with default parameters. The rates are estimated between two consecutive time points, therefore the sample from 8h was not included because it is missing the corresponding MT data. We filtered the results to identify the changing point using a score threshold of 0.9 and a FDR equal to 0.05.

