## Supplementary material for "Integration of absolute multi-omics reveals translational and metabolic interplay in mixed-kingdom microbiomes": Summary of supplementary figures and tables

This PDF file includes:

I. Supplementary figures and legends
II. Supplementary tables and legends

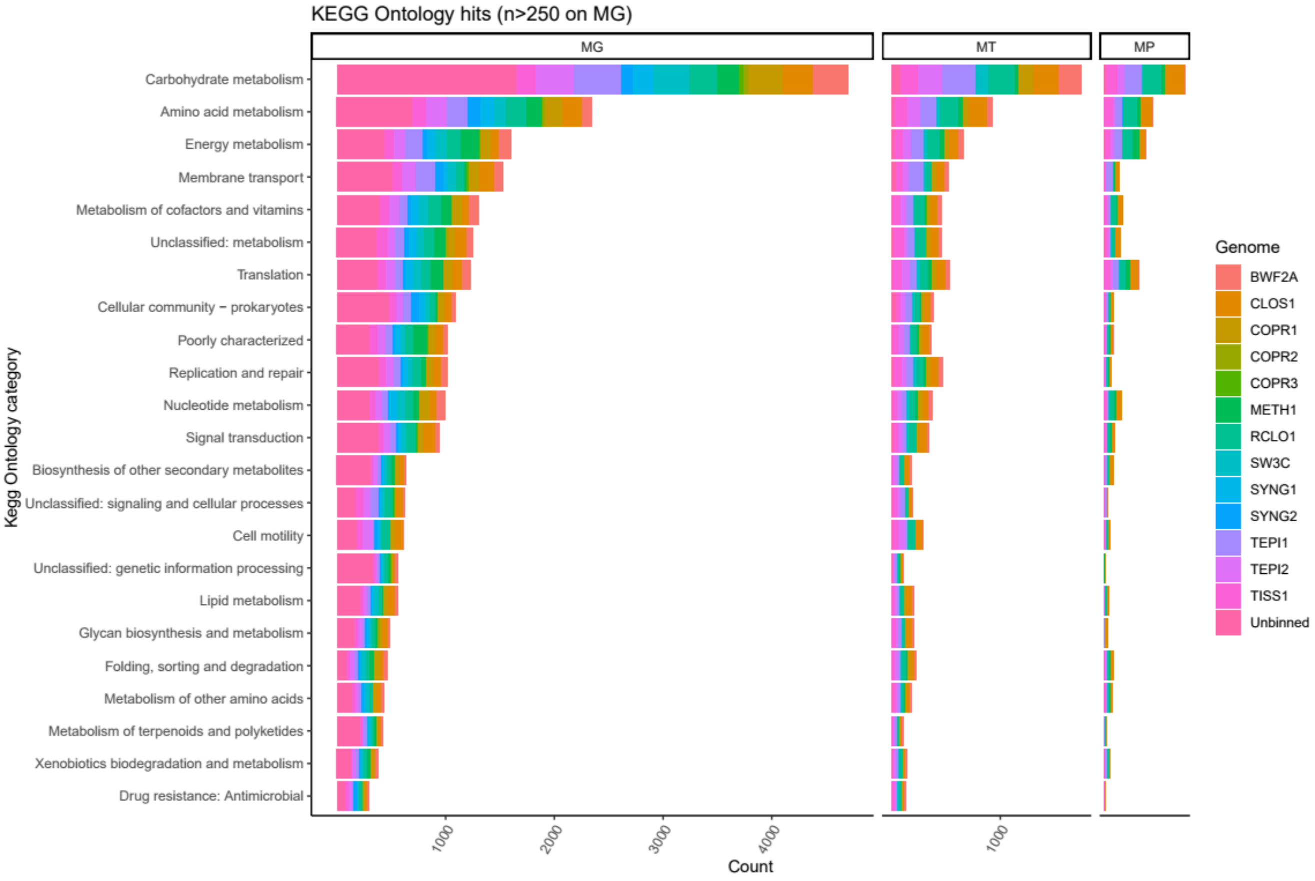


**Supplementary figure 1.** Counts of KEGG Orthology hits in the different omic layers per KEGG category. The colors represent the metagenomic bins to which the annotated ORFs belong. The sorting of the KEGG categories was performed using the counts for the MG (hence relative to the genomic potential), and it is more conserved in the MT (Kendall tau: 0.77, $p<10^{-8}$) than in the MP (tau 0.68, $p<10^{-5}$).

**Supplementary table 1.** Map among ORF name, contig and bin.

**Supplementary table 2**. Map between ORF name and KEGG Orthology code.

**Supplementary table 3**. Filtered expression table for MT, measures in log10(molecules+1). Each column corresponds to an ORF, each row to a sample.

**Supplementary table 4**. Filtered expression table for MP, measures in log10(molecules+1). Each column corresponds to an ORF, each row to a sample.

**Supplementary table 5**. The main output from the peca analysis. Each row repesents an ORF, the colums called R<number> and D<number> contain the estimate of the translation and protein degradation rates respectively. The CPS<number> columns contain the score for being a change point in the parameters and the FDR<number> columns store the False Discovery Rate associated to the CPS.
